# An allelic resolution gene atlas for tetraploid potato provides insights into tuberization and stress resilience

**DOI:** 10.1101/2025.06.26.661617

**Authors:** Julia Brose, Dionne Martin, Yi-Wen Wang, Joshua C Wood, Brieanne Vaillancourt, John P. Hamilton, C. Robin Buell

## Abstract

Tubers are modified underground stems that enable asexual, clonal reproduction and serve as a mechanism for overwintering and avoidance of herbivory. Tubers are wide-spread across angiosperms with some species such as *Solanum tuberosum* L. (potato) serving as a vital crop for human consumption. Genes responsible for tuber initiation and disease resistance have been characterized in potato including *StSP6A*, a homolog of Flowering Time, that functions as tuberigen, the equivalent of florigen. To elucidate additional molecular and genetic mechanisms underlying potato biology including tuber initiation, tuber development, and stress responses, we generated a developmental and abiotic/biotic-stress gene expression atlas from 34 tissues and treatments of Atlantic, a tetraploid cultivar. Using the haplotype-phased tetraploid Atlantic genome assembly and expression abundances of 129,218 genes, we constructed gene coexpression modules that represent networks associated with distinct developmental stages as well as stress responses. Functional annotations were given to modules and used to identify genes involved in tuberization and stress resilience. Structural variation from a pan-genomic analysis across four cultivated potato genome assemblies as well as domestication and wild introgression data allowed for deeper insights into the modules to identify key genes involved in tuberization and stress responses. This study underscores the importance of transcriptional regulation in tuberization and provides a comprehensive framework for future research on potato development and improvement.

## Introduction

Tubers are modified underground stems that serve as a mode of asexual reproduction with some species cultivated as a food source for human consumption. Tubers can be found throughout the Angiosperms, some of which, are important food crops such as potato, oca, yam, and mashua. Potato (*Solanum tuberosum* L.), a model for tuber development, is the fourth most important crop based on production value in the world, worth approximately US$120 billion (2022, http://www.fao.org/faostat/). Potatoes rely on production from seed tubers rather than true seed. Tuber formation is a complex developmental process that involves multiple pathways including genes involved in hormone regulation (*StPIN1, StLOG1*), photoperiod regulation (*StCO*), maturity (*StCDF1*), as well as carbohydrate transport (*StSUT1, StSWEET11)* (Navarro *et al*., 2011; Abelenda *et al*., 2016; Abelenda *et al*., 2014). The key tuberigen protein StSP6A is a mobile protein that travels from the leaf through the phloem into the stolon, an underground stem that initiates tuber development (Navarro *et al*., 2011; Tang *et al*., 2022; Nicolas *et al*., 2022). The first stage in tuber development is the formation of a hooked stolon which then undergoes cell division to form a swollen stolon before finally developing into a tuber.

Cultivated potato is an autotetraploid with a highly heterozygous genome (Hoopes *et al*., 2022). Due to its clonal reproduction, potato has a high genetic load as retained deleterious and dysfunctional alleles are not purged through meiosis. Furthermore, intragenomic structural variation has been reported that results in allele dosage variation across the genome. In addition, potato has substantial wild species introgressions, some of which are associated with adaptation to stress response and disease resistance (Hoopes *et al*., 2022). Studies on domestication and improvement examining both cultivars and landraces revealed selection on genes in key pathways associated with tuberization, carbohydrate metabolism, glycoalkaloid biosynthesis, circadian rhythm, endoreduplication, and fertility (Hardigan *et al*., 2017).

Genome assemblies of haplotype-resolved chromosome-scale tetraploid cultivars include: North American cultivars such as Atlantic and Castle Russet, the Asian cultivar Cooperation 88, and the European cultivar Otava (Sun *et al*., 2022; Hoopes *et al*., 2022; Bao *et al*., 2022). However, despite the economic importance of potato, an extensive developmental gene expression atlas for potato is not available. This study reports on an allele-resolved gene expression atlas in Atlantic that incorporates a tuber developmental series, sprouting potato, leaf diurnal time course, leaf abiotic and biotic stresses, stem, root, root abiotic and biotic stresses, two stages of floral development, and two stages of fruit development. Allele-resolved gene expression abundances were then used to construct gene coexpression networks to facilitate identification of genes involved in key potato biological processes.

## Results and Discussion

### An allelic-resolved gene expression atlas for potato

To understand potato development and responses to stress, we generated a replicated gene expression atlas using RNA-sequencing (RNA-seq) of the tetraploid potato cultivar Atlantic. In total, 34 tissues and treatments were harvested including seven stages of tuber development from hooked stolon to mature tuber (Tuber Stage 5; Figure 1a), a 24-hour diurnal leaf time course, stem, flower (closed and open flowers), fruit (immature, mature), sprout, and eight abiotic/biotic stress treated samples (leaves and roots). Allelic expression abundances (transcripts per million (TPM)) were calculated using the phased, haplotype-resolved chromosome-scale Atlantic genome sequence (Hoopes *et al*., 2022). Technical replicates with a Pearson’s correlation coefficient of less than 0.95 were flagged as outliers and removed resulting in a total of 95 RNA-seq libraries. Expression abundances of replicates were averaged resulting in expression data for 34 tissues and treatments; of 133,989 total genes in the Atlantic genome, 130,927 were expressed (TPM > 0) in at least one tissue or treatment. Principal component analysis (PCA) revealed distinct groupings of above- and below-ground organs along the Principal Component 1 accounting for 37.6% of variance (Figure 1b). As expected, all stages of stolon and tubers grouped together indicating that these tissues are more similar to each other than the other tissues. The second largest contribution to variance amongst the samples was the separation between the two flowering stages (closed vs open flowers) indicating that these stages are distinct. Hierarchical clustering also revealed clear separation between above-ground organs, floral/fruit structures, and below-ground organs (Figure 1c). There was distinct grouping of Tuber Stages 1, 2, 3, 4, and 5 that were differentiated based on size, whereas Hooked Stolon and Swollen Stolon clustered with Sprouts, reflective of their stem-derived meristematic state. Although stem clustered with leaves, stem has high correlation values (∼0.8) with the stem-derived underground structures, stolons and tubers.

**Figure 1.**
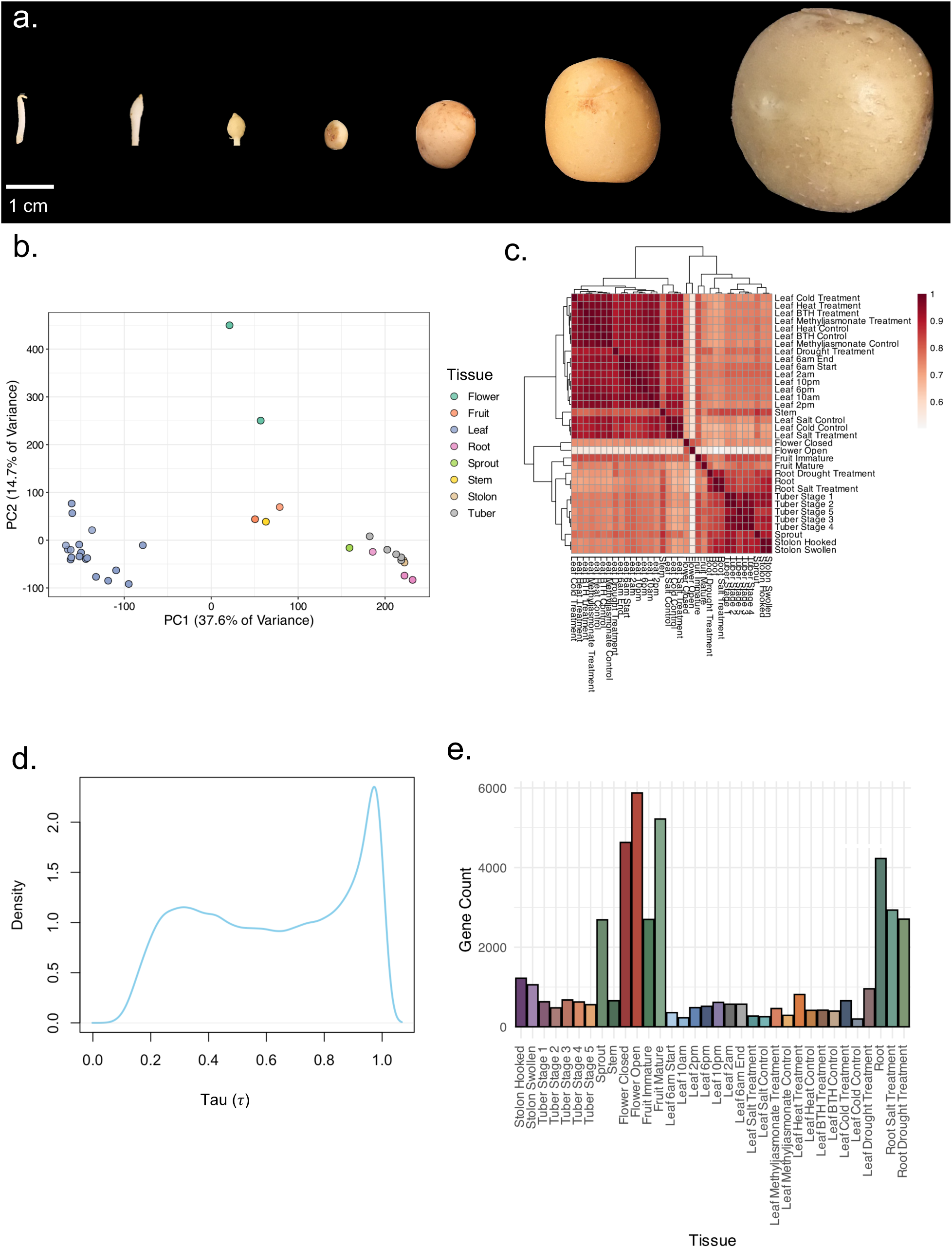
Tuber Developmental Series and RNA-Seq Data Analysis from Tissue Atlas. A. Photos of tuber developmental tissue series. From left to right, hooked stolon, swollen stolon, tuber stage 1, tuber stage 2, tuber stage 3, tuber stage 4, and tuber stage 5. B. PCA of Atlantic RNA-Seq data. Color represents the broad tissue type: flower, fruit, leaf, root, sprout, stem, stolon, or tuber. C. Hierarchical clustering of Atlantic RNA-Seq data. Colors indicate the correlation coefficient (r) value between the various tissue types where dark red represents a value closer to 1 and white closer to 0. D. Gene tissue specificity density plot. A density plot of *τ* values generated for all alleles. E. Peak expression of tissue-specific genes. Tissue-specific genes were determined by a *τ* value greater than or equal to 0.8. The number of genes with peak expression is plotted by the tissue type.

Preferential allele expression occurs when one allele is expressed more abundantly than the other alleles and has been described previously in tetraploid potato which is attributable to the high degree of heterozygosity and structural variation among the four alleles (Pham et al., 2017; Hoopes et al., 2022). However, these studies were limited to a single leaf and tuber sample. With access to a robust gene atlas for tetraploid potato, we further explored the extent of preferential allelic expression. In total, 31,156 allelic groups were found within Atlantic based on their synteny with the doubled monoploid reference genome DM (Pham *et al*., 2017). As expected, based on the ploidy, the average number of alleles in each group is 2.97 with 14,238 found in group 0 (not present in an allelic group), 24,459 in an allele group of 1 (present in 1 copy in Atlantic, mono-allelic), 27,005 in an allele group of 2 (present in 2 copies in Atlantic, bi-allelic), 27,927 in an allele group of 3 (present in 3 copies in Atlantic, tri-allelic), and 25,050 in an allele group of 4 (present in 4 copies in Atlantic, tetra-allelic). We examined genes across the 34 different tissues/treatments and found that 14,335 allele groups (46% of allele groups) displayed preferential gene expression. The mean number of tissues/conditions in which they are preferentially expressed was 7.15 with a total preferential allele expression observed across 94,390 across all tissues/conditions. Salt leaf showed the highest number of genes with preferential allele expression with 5,319 alleles whereas only 331 alleles are preferentially expressed in the 6am Leaf.

Tissue and treatment specificity of gene expression across the samples was measured using the Tau (*τ*) value (Kryuchkova-Mostacci and Robinson-Rechavi, 2017). This value ranges from 0 to 1 in which globally expressed genes have a value of 0 and genes with tissue- or treatment-specific expression have a value closest to 1; genes with *τ* values greater than or equal to 0.8 were considered tissue-specific (Figure 1d). Of the 130,927 expressed genes in the Atlantic genome, 45,279 (34.6%) are tissue/treatment-specific which were then categorized based on their peak expression to gain further insights into the distribution of expression among tissue or treatment (Figure 1E). Unsurprisingly, tissue types with the largest number of tissue-specific genes included those with smaller sampling sizes (sprout, flower, fruit, root) but excluded stems. These four tissues had less sampling across development, time, and treatments and represent distinct organs from the leaf, stolon, and tuber sets that were sampled across development and/or stresses. Unlike these four tissues, stem had a large number of tissue/treatment-specific genes, suggesting that while stolon is a stem-derived structure, it is distinct from aboveground photosynthetic stems.

The gene expression atlas also included four abiotic (cold, drought, heat, salt) and two biotic (benzothiadiazole (BTH), methyl jasmonate (MeJa)) stress-treatments of leaves and two abiotic stress-treatments (drought and salt) of roots. BTH is known to induce a systemic resistance response to a broad range of biotic stressors across many plant species (Bokshi *et al*., 2003). MeJa treatment mimics herbivory and elicits a defense response from plants (Wu *et al*., 2008). Differentially expressed genes (DEGs) were identified between stress and control tissues using a p-value < 0.05 and logfold change > |2|. In total, 39,039 unique DEGs were identified across all stress conditions and tissues. The treatment with the largest number of upregulated DEGs is drought leaf with 11,373 genes (Table 1) whereas the lowest number of upregulated DEGs, 874, was salt-treated root (Table 1). The number of DEGs from abiotic stress across all tissues and treatments was greater than biotic stress treatments. There is a small overlap between the conditions with 756 DEGs downregulated and 1,552 upregulated genes between all stresses.

**Table 1.** Differentially Expressed Genes in Abiotic and Biotic Stress Treatments.

|  | Treatment | Up Regulated | Down Regulated |
| --- | --- | --- | --- |
| Abiotic Stress | Cold Leaf | 9,134 | 4,091 |
|  | Drought Leaf | 11,373 | 7,439 |
|  | Drought Root | 4,910 | 8,293 |
|  | Heat Leaf | 2,391 | 1,693 |
|  | Salt Leaf | 1,405 | 3,244 |
|  | Salt Root | 874 | 1,249 |
| Biotic Stress | BTH Leaf | 2,251 | 794 |
|  | MeJa Leaf | 1,565 | 1,411 |

### Expression of genes implicated in tuber initiation, growth, and development and in response to stress

In potato, multiple genes involved in tuber initiation, growth, and development have been characterized, yet most gene expression studies are limited to a few tissues or timepoints and have primarily used quantitative reverse transcription polymerase chain reaction for quantification. Thus, our 34 tissues/treatments gene expression atlas provides an expansive dataset to not only characterize gene expression across development and stress of these known genes but also to link new genes with putative functions. Using a set of 57 genes (212 alleles) previously implicated in tuberization, growth and/or development, we characterized their expression in our 34 tissue/treatment replicated atlas. Genes were binned into four main functional categories: Hormone response, Tuber initiation, Tuber inhibition, and Starch/carbohydrate metabolism (Figure 2). A total of 16 genes (32 alleles), *StBEL30*, *StBRC1b/IT1*, *StCO1*, *StCO2*, *StGA20ox1*, *StGA2ox1*, *StPHYB2*, *StPIN1*, *StPOTH1*, *StPURINE TRANSPORTER 2*, *StSWEET10A, StSWEET17B, StSWEET1C, and StSWEET1F* exhibited tissue-specific expression based on a Tau value greater than 0.8. The transcription factor, *StBRC1b/IT1* (all 3 alleles), which has been shown to function in tuber initiation exhibited tissue specific expression (tau values: 0.88, 0.88, 0.87) with peak expression at the hooked stolon stage. *StPOTH1* has also been shown to function in tuber initiation with 3 of 4 alleles exhibiting tissue specific expression (tau values: 0.87,0.82,0.81) with peak expression in the sprouting tuber as well as high expression in the hooked stolon. Genes with leaf specificity include *StCO1* (all 3 alleles; tau values: 0.92, 0.88, 0.87) and *StCO2* (1 of 4 alleles; tau value: 0.82). Both *StCO1* and *StCO2* gene expression follow a diurnal rhythm with peak expression during early morning hours. *StSP6A* is expressed in leaf tissue and the StSP6A protein moves from the leaf into the stem and then into the hooked stolon to signal tuberization (Navarro *et al*., 2011). The StSP6A protein, also known as tuberigen, forms an activation complex with St14-3-3, and StAST1 (Sun *et al*., 2024). Interestingly, both *St14-3-3* and *StAST1* are expressed in most other tissues whereas, *StAST1* has higher expression in early tuberization (hooked and swollen stolon). The alternative tuberigen complex relies on StABL1 rather than StAST1 which is also constitutively expressed (Sun *et al*., 2024). Therefore, tuber initiation relies on the StSP6A protein traveling from the leaf to the underground stem structure interacting with binding partners that are not selectively expressed.

**Figure 2.**
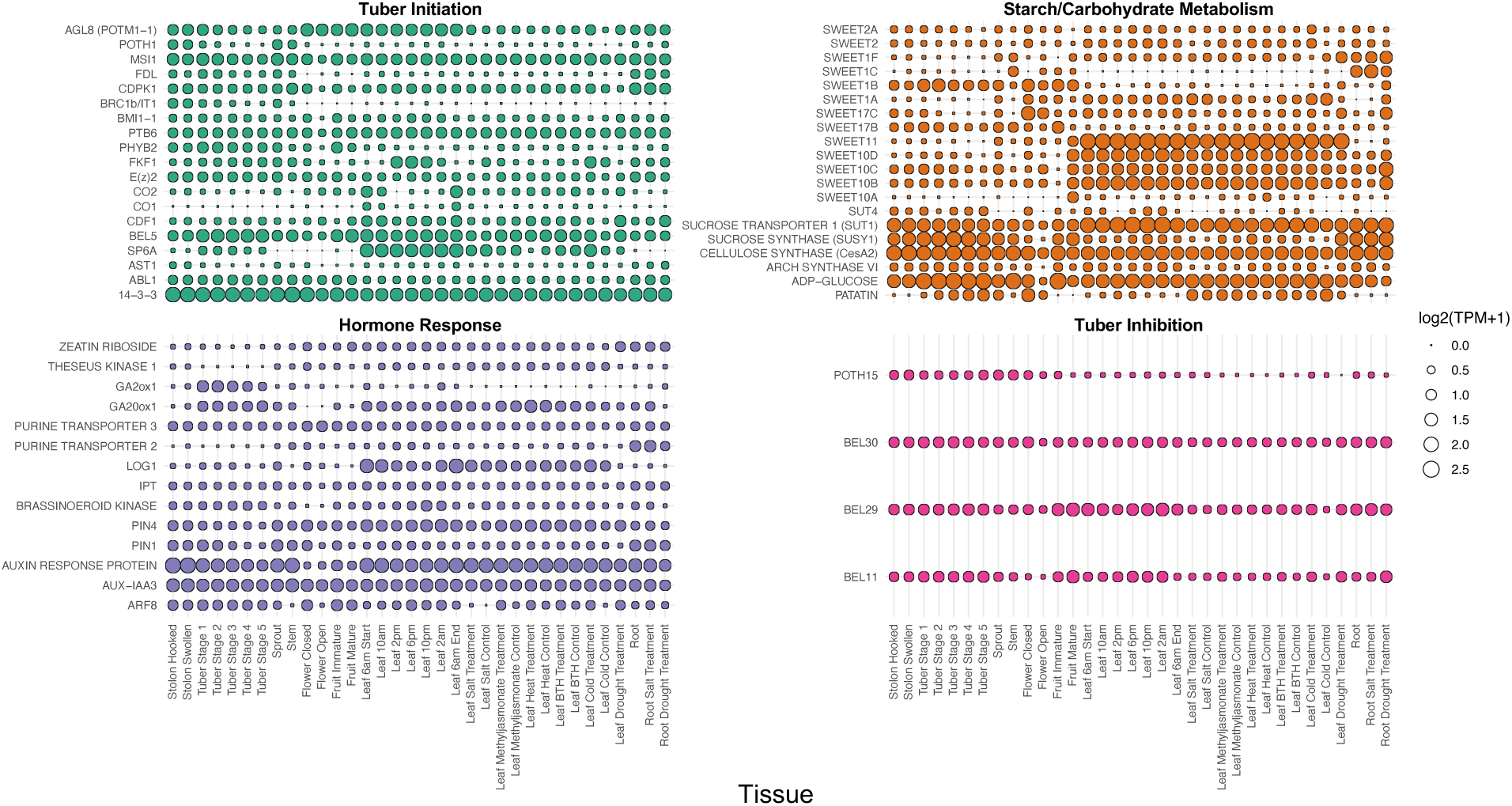
Expression of Tuberization Genes. A representative allele of known tuberization genes expression values (log2(TPM+1)) are plotted by the 34 tissues collected. The plot is broken into different stages of tuberization including hormone response, tuber initiation, tuber inhibition, and starch/carbohydrate metabolism. The allele with the highest overall expression was chosen as the representative example.

Throughout the process of tuber development and maturation (hooked stolon to Tuber Stage 5), expression of *StPOTH1, StBRC1b, StPIN1,* and *StPIN4* decrease while other genes*, StGA2ox1*, *StGA20ox1*, and *PATATIN* increase. Not surprisingly, genes involved in starch, carbohydrate, and glycoprotein biosynthesis were expressed across multiple tissue types (Figure 2). Patatin, a storage glycoprotein also known as tuberin, is vital in the composition of the potato tuber. Approximately 40% of soluble proteins in the potato are from patatin (Liu *et al*., 2003) consistent with its expression increasing 4-fold from Tuber Stage 1 until Tuber Stage 5. Patatin is also known for its anti-fungal activity to *Phytophthora infestans,* and inhibition of *Meligethes* spp. and *Diabrotica* spp. larval growth (Ralet and Guéguen, 2000; Creusot *et al*., 2011; Sharma *et al*., 2004; Åhman and Melander, 2003; Strickland *et al*., 1995). Patatin was also expressed in leaf and flower tissues but not fruit.

The set of 57 tuber initiation, growth and development genes (212 alleles) were also examined for expression changes in response to abiotic and biotic stressors. We found that 39 known genes (190 alleles) involved in tuberization are differentially expressed during abiotic and biotic stress tissues with 30 of the known genes (121 alleles) upregulated during abiotic/biotic stress treatments. The stressed tissue was collected from 4-week old plants which are initiating stolons but had yet to start to develop tubers; thus, we anticipate that the earliest stages of stolon formation could be affected by the stress conditions. All alleles of *StARF8* were differentially expressed under at least one condition. In particular, *StARF8*e was upregulated in four of the eight stress treatments (BTH leaf (log2fold: 7.27, p-value: 7.26E-4), MeJa leaf (log2fold: 4.55, p-value: 9.62E-5), Cold leaf (log2fold: 5.25, p-value: 2.02E-2), Drought leaf (log2fold: 6.68, p-value: 4.68E-5)). In *Arabidopsis thaliana, AtARF8* was indicated as a key gene in regulating auxin responses to abiotic and biotic stress (Truskina *et al*., 2021). In Atlantic, the expression pattern of *StARF8*e suggests a similar role. Other functions of *StARF8* in potato include changes in brassinosteroid levels though the activation of the transcription factor *DWARF4* to direct cell wall formation and leaf shape (Xiong *et al*., 2021). Additionally there are changes to the developing underground stem due to *StARF8* (Bao *et al*., 2025; Kondhare *et al*., 2021). Unlike *ARF8e,* the remaining alleles are differentially expressed in fewer tissues indicating potential alternate roles of the alleles. For example, *StARF8a* is only differentially expressed in the leaf during cold treatment (log2fold: 3.34, p-value: 0.02) indicating the lack of auxin response during other abiotic and biotic stress treatments. Additionally, *StARF8b* (log2fold:-2.95, p-value: 0.024) and *StARF8d* (log2fold: -3.69, p-value: 0.010) are downregulated in the root under drought treatment. These findings suggest that StARF8e may play a central role in mediating auxin-driven responses to environmental stress, while the differential expression of other *ARF8* alleles hints at functional diversification in stress signaling pathways. It also highlights the power of allelic resolution of gene expression obtained through the phased, haplotype-resolved Atlantic genome assembly and allelic-resolved gene expression profiles.

### Gene coexpression modules implicate new genes involved in potato biology

Our allele-resolved gene expression atlas was used to construct a gene coexpression network resulting in 26 modules containing 129,218 of the 130,927 genes that could be associated with specific developmental tissues, stages, and/or treatments (Figure 3a). The modules varied in size from 10,950 genes to 20 genes (Figure 3a); not surprisingly, preferential allele expression was observed in the gene coexpression modules with 70% of genes with at least two alleles placed in different modules indicating divergent expression patterns across alleles. Initial efforts to annotate modules with respect to biological function using all genes within the module was uninformative due to the under annotation of genes in the Gene Ontology (GO) database that are specific to potato. Thus, to assign modules to function, hub genes were first defined as the top 5% of genes in the modules based on the number of edges. Next, peak average gene expression for each module was used in combination with hub gene GO term enrichment tests to assign a biological function for each module. From the resolved modules, we identified new genes involved in tuber initiation, growth, development and stress responses.

**Figure 3.**
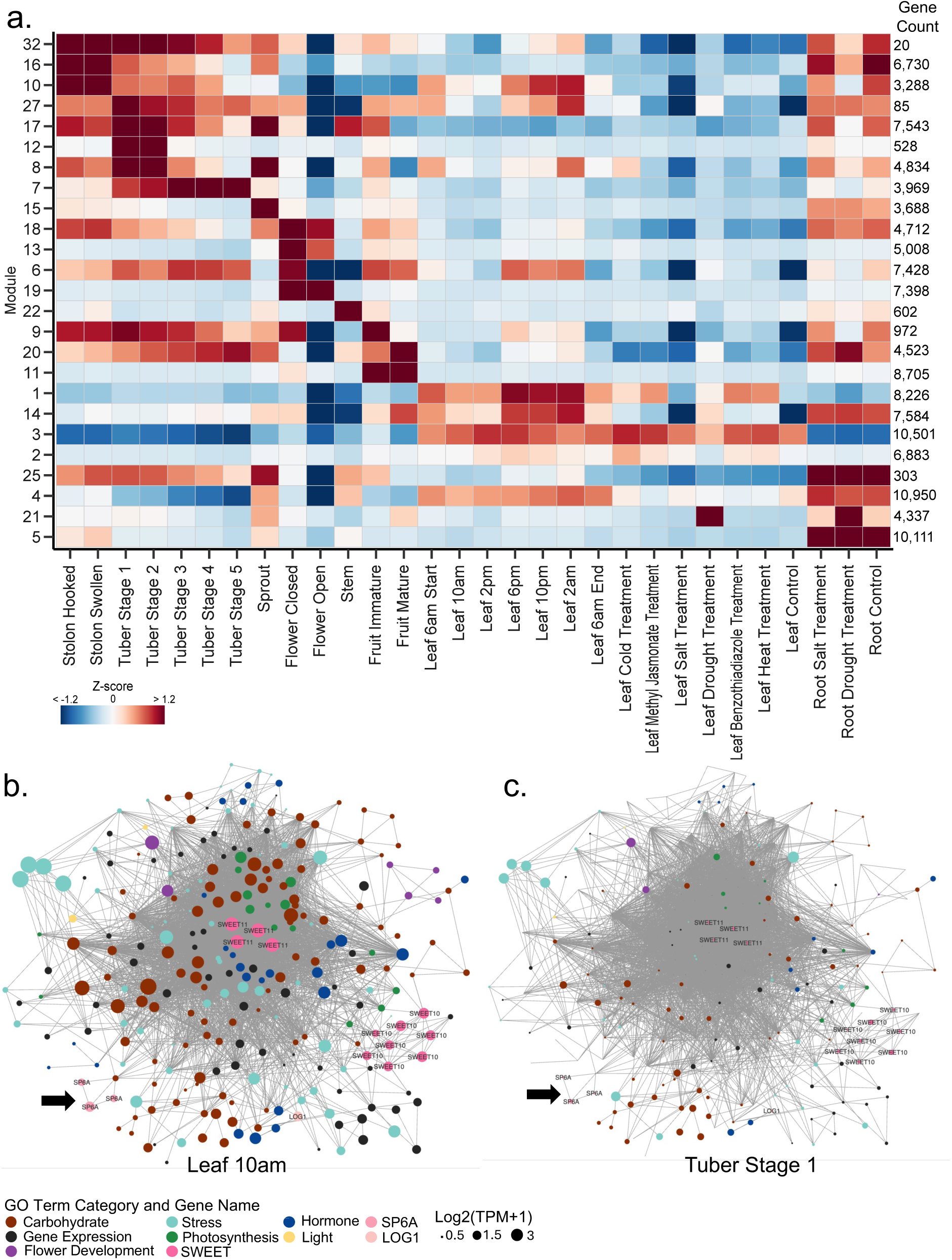
Gene Coexpression Modules and *StSP6A* Submodule Networks. a. Average gene coexpression module expression. The expression across tissues is represented by the Z-score with high values ( > 1.2) in red and low (< -1.2) in blue. The colors along the x-axis correspond to the tissue type. The numbers along the right side indicate the number of genes in the module. b-c. *StSP6A* submodule network. Submodule 5_2 containing *StSP6A* (indicated by black arrow) where the size of the circle represents the expression for the leaf 10am timepoint (b) or tuber stage 1 (c). The color represents the GO term annotation.

#### SP6A: The tuberigen signal

To identify gene networks involved in tuber development, we began with a curated list of 57 genes (218 alleles) implicated in tuber initiation, growth and development. Of these, 212 alleles were represented across 21 of the 25 gene coexpression modules. Of the 218 alleles, nine alleles are hub genes: *StBEL14* (1 allele, Module 1), *StCO1* (2 alleles, Module 2), *StSWEET11* (4 alleles, Module 3), *StPIN1* (1 allele, Module 15), and *St*E(z)2 (1 allele, Module 18). To further understand genes involved in early tuberization signaling we investigated the tuberigen gene *StSP6A,* an ortholog of *FLOWERING LOCUS T* in *A. thaliana* (Navarro *et al*., 2011). All alleles of *StSP6A* were present in Module 3, a leaf-exclusive module with no expression in any underground stem tissues. Module 3 contains 10,501 genes. To refine the network and identify genes most closely coexpressed with *StSP6A*, we generated submodules. *StSP6A* is present in Submodule 3_2 which also contains the known tuberization genes, *StLOG1, StSWEET10B, StSWEET10D,* and *StSWEET11* (Kondhare *et al*., 2021).

Submodule 3_2 contains 1,625 genes (3,571 alleles) and is enriched in GO terms associated with “generation of precursor metabolites and energy” (GO:0006091, p-value: < 1e-30), “response to light stimulus” (GO: 0009416, p-value: < 1e-30), and “photosynthesis” (GO:0015979, p-value: < 1e-30). Floral development genes are particularly interesting due to the cooption of *FLOWERING LOCUS T* as the key tuberization signal in potato and Submodule 3_2 contains genes that encode orthologs of genes associated with flower development based on their GO term annotation including an ortholog of *HEADING DATE REPRESSOR 1* (HDR1,1 allele), SQUAMOSA promoter binding protein (SBP, 4 alleles), and *CO-like* transcription factor (2 alleles). The *CO-like* transcription factor is of interest due to the known activity of *StCO* in the regulation of tuberization (Abelenda *et al*., 2016). *HEADING DATE REPRESSOR 1* represses the expression of *HEADING DATE 1* which is a known ortholog of *CO* and involved in the photoperiod control of flowering in rice (Sun *et al*., 2016). *SBP* and their orthologs are known to be involved in floral development, regulation of GA signaling, fruit ripening, and plant architecture in maize, rice, and *A. thaliana* (Chen *et al*., 2010). These genes could be additional genes in floral signaling that have been coopted for tuber development.

We examined the network topology and composition of Submodule 3_2. In Figure 3b, the size of the nodes indicates expression abundance in the leaf at 10am. Surprisingly, alleles of *StSP6A* are not the most highly connected genes within the network with 1,855 edges connected to *StSP6Aa/c* (52% of the module) to 1,952 edges for *StSP6Ab* (55% of the module). Rather, the sucrose transporter *StSWEET11* is more highly connected are connected within the network with *StSWEET11b* having 3,290 edges (92% of module) and *StSWEET11c* having 3,265 edges (91% of module). This includes genes involved in floral development, photosynthesis, and carbohydrate transport (Figure 3b). Central to the network are four alleles of *StSWEET11,* known to be post-translationally regulated by *StSP6A* (Abelenda *et al*., 2019). In addition to *StSWEET11,* four alleles of *StSWEET10B,* and four alleles of *StSWEET10D* are present within Submodule 3_2 bringing the total number of genes involved in carbohydrate metabolism within this submodule to 154.

To understand changes in source-sink balance as well as preferential allele expression during the transition into underground stem production, we examined expression levels of genes within Submodule 3_2 at Tuber Stage 1, the first stage of tuber bulking, relative to their expression at Leaf 10 AM. Figure 3c shows the same submodule for *SP6A,* but with nodes sizes determined by expression at Tuber Stage 1. One allele, *StSP6Ab,* is expressed more abundantly than the other two alleles highlighting preferential allelic expression (Figure 3c). There are also three other groups of abundantly expressed genes during Tuber Stage 1. The four alleles of *HEVEIN-LIKE, pathogenesis-related 4* (light blue-stress nodes), two alleles of *StCO-like 4* (purple-flower development), and a single allele of *ascorbate peroxidase 3* (light blue-stress).

#### The hooked stolon: Site of tuber initiation

With their high expression in the hooked stolon tip and swollen stolon tip (Figure 3a), *StPOTH1b, StPOTH1c,* and all *StBRC1b* alleles were placed in Module 16, a module with peak expression in the hooked stolon stage. To identify additional genes involved in tuber initiation and to resolve co-regulation relationships within our coexpression networks, we identified modules involved in tuber initiation based on peak expression in the hooked and swollen stolons with limited expression in other tissues as well as the presence of known, characterized tuberization genes within the module. Three modules (Modules 32, 16, and 10) containing 20, 6,730, and 3,288 alleles, respectively, met these criteria. Gene ontology (GO) term enrichment for molecular function in Module 16 included “DNA-binding transcription factor activity” (GO:0003700, p-value 2.10E-62) and “transcription regulator activity” (GO:0140110, p-value 4.52E-52) suggesting this module was involved in regulation of tuber initiation in the stolon. Module 32 was enriched for biological functions involving the brassinosteroid hormone including: “cellular response to brassinosteroid stimulus” (GO:0071367, p-value 1.01E-4), “brassinosteroid mediated signaling pathway” (GO:0009742, p-value 1.01E-4), and “response to brassinosteroid” (GO:0009741, p-value 1.01E-4) suggesting genes involved in hormone responses. Module 10 was enriched in biological functions associated with protein modification (GO: GO:0036211, p-value 5.04E-39) and protein phosphorylation (GO: GO:0006468, p-value 1.10E-37).

From the GO enrichment analysis of our gene coexpression modules, we hypothesized that Module 16 contained key regulatory factors associated with initiation and transition of the stolon to a tuber including the transcription factors, *StPOTH1* and *StBRC1b/IT1*, previously shown to be involved in tuber initiation (Chen *et al*., 2004; Tang *et al*., 2022). Approximately 10% of Module 16 (6,730 genes) are transcription factors representing 39 transcription factor families. To refine coexpression relationships in Module 16, we constructed submodules to further separate genes with near exclusive expression in the hooked and swollen stolon relative to other tissues or treatments. Submodule 16_3 contains 2,088 genes including *StPOTH1* and *StBRC1b* alleles with high expression in hooked and swollen stolons and limited expression in sprout and root tissue. Gene ontology term enrichment of Submodule 16_3 revealed the most significant GO term was “DNA-binding transcription factor activity” (GO:0003700; p-value 1.06E-10) suggesting that this submodule represents a network of key regulatory genes that function in tuber initiation and development. Submodule 16_3 contains 158 total transcription factors from 27 different transcription factor families and is significantly enriched in HB-WOX (p-value 7.53E-04) and HB-other (p-value 8.41E-03) transcription factors. To dissect gene regulation during tuber initiation, we constructed a transcription factor coexpression network for Submodule 16_3 (Figure 4) highlighting not only the complex connectivity between transcription factors but also the dynamic levels of gene expression with the MYB and AP2/ERF-ERF families having the highest expression.

**Figure 4.**
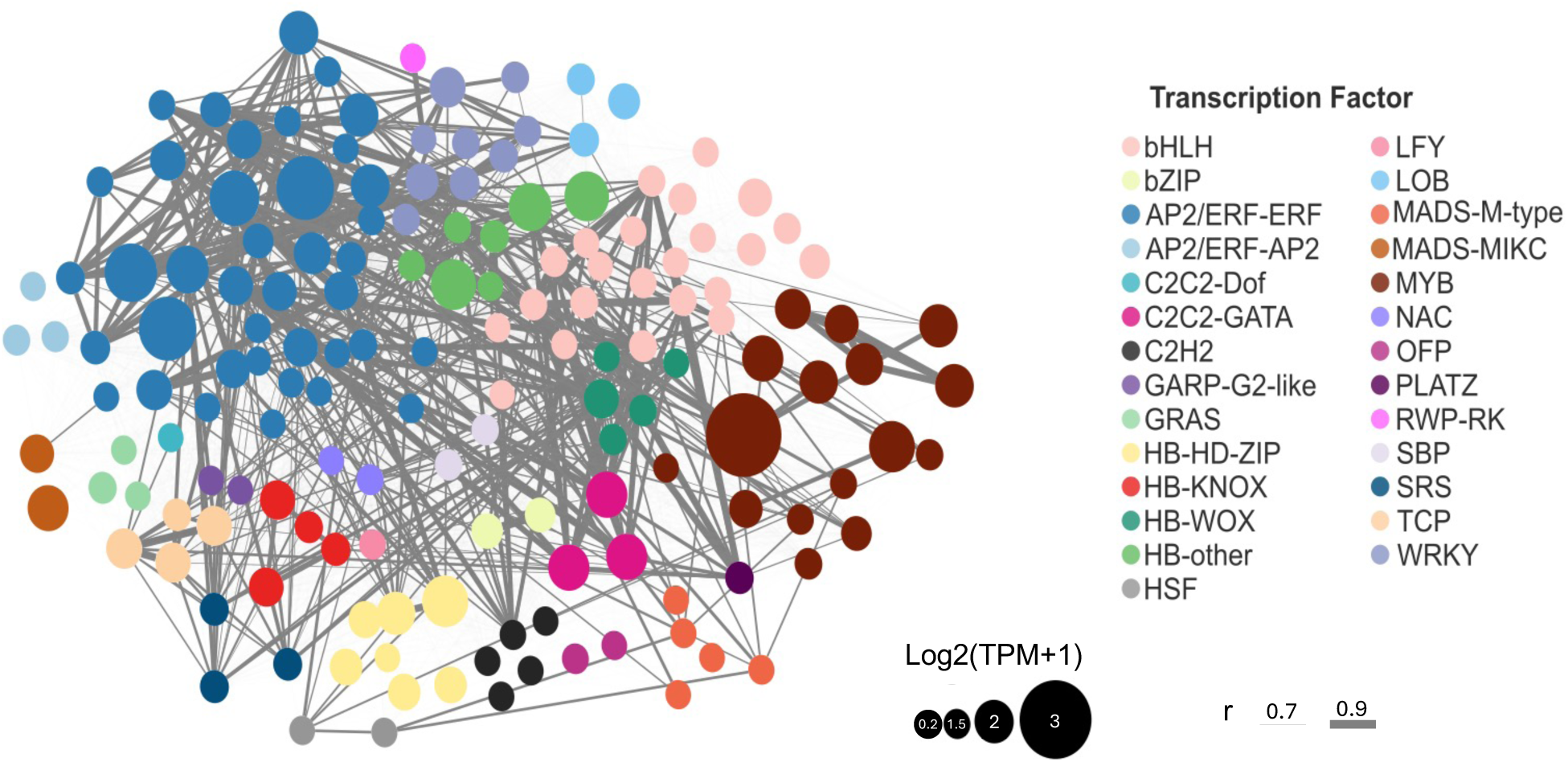
Submodule 16_3 Transcription Factor Network. Weight of the edges is determined by the correlation value between nodes. Size of the node is determined by the log2(TPM+1) expression abundance during the hooked stolon stage.

#### Transition from a swollen stolon to a bulking tuber

We investigated the earliest stages of tuber formation by examining Module 12 with peak expression during Tuber Stage 1. At Tuber Stage 1 there is expansion of the swollen stolon and transition into a tuber. There still may be a hook present yet the tuber is not completely rounded (Figure 1a). Module 12 (707 total genes) captures this developmental stage and has peak expression in Tuber Stage 1; hub gene GO term enrichment suggests genes that function in catalytic activity and cellular homeostasis. The transition into bulking relies on genes involved in cell wall modification and both pectate lyase 2 and endo-1,4-beta-xylanase 5 are hub genes in Module 12. Pectate lyases have been implicated in tissue development, petal abscission, lateral root emergence, leaf senescence, and pollen development (Zheng *et al*., 2018). Additionally, a hub gene within Module 12 encodes a *COBRA-like protein-7 precursor (COBL7)*. In *A. thaliana, COBL7* regulates cellulose deposits required for stomata development (Ge *et al*., 2024). As a modified stem, tubers also develop lenticels in the potato skin to facilitate the exchange of oxygen and carbon dioxide analogous to stomata (Bethke, 2023) suggesting that this gene may facilitate the development of lenticles during tuber development.

### Structural variants

Structural variation has been reported in several crop genomes including potato in which intra-genome and inter-genome structural variation occurs (Hoopes *et al*., 2022; Hardigan *et al*., 2017). Pangenomic analysis in peanut revealed structural variants in exon and promoter regions affect gene expression directly relating to yield through genes involved in seed size and weight (Zhao *et al*., 2025). Similar pangenomic analysis have been occurred in cucumber where it was found that duplicated genes tended to be involved in reproduction morphology whereas deleted regions were involved in abiotic stress and histone modification (Zhang *et al*., 2015). In potato, structural variants have been associated with disease resistance (Lian *et al*., 2025). Based on representation in a syntenic pan-genome, genes can be defined as core, soft core, shell, or cloud based on presence in all (or nearly all) accessions, a subset of accessions, or private to one or two accessions (Gordon *et al*., 2017). With access to four haplotype-resolved chromosome-scale genome assemblies of tetraploid cultivated potato (Hoopes *et al*., 2022; Bao *et al*., 2022; Sun *et al*., 2022), we performed a pan-genome analysis using a synteny-based approach which is much more stringent compared to orthology based pan-genomes that do not consider gene order conservation. In total, the four tetraploid assemblies representing 16 haplotypes and contained 455,849 syntenic genes revealing both shared evolutionary origin and genomic position and were then binned into core, soft core, shell, cloud and private genes. Core genes 30,560 genes (6%) are central to cultivated potato and were required to be present in all 16 haplotypes which represents. Soft core genes are shared between 12-15 haplotypes with 199,704 genes (39%). Shell genes are between 2-11 haplotypes with 238,719 genes (46%) and cloud genes which are unique to a single haplotype with 46,962 genes (9%). Within Atlantic, 90,244 genes were placed into syntenic groups (Figure 5a) with a total of 7,123 core, 42,761 soft core, 34,816 shell, 5,544 cloud, and 40,681 genes that were private to Atlantic. We examined the pan-genomic context of the Atlantic genes to understand how structural variation impacted coexpression network especially connectivity to other genes within the module. From our coexpression analyses, there are 6,722 hub genes (defined as the top 5% of highly connected genes within the modules) within the 25 modules. Surprisingly, core genes represent only 6.4% of total hub genes in the coexpression modules whereas soft core genes represent 39.3%, shell genes represent 23.5%, cloud genes represent 1.8% of all hub genes, and 29% of hub genes are unplaced in a pangenomic context (private).

**Figure 5.**
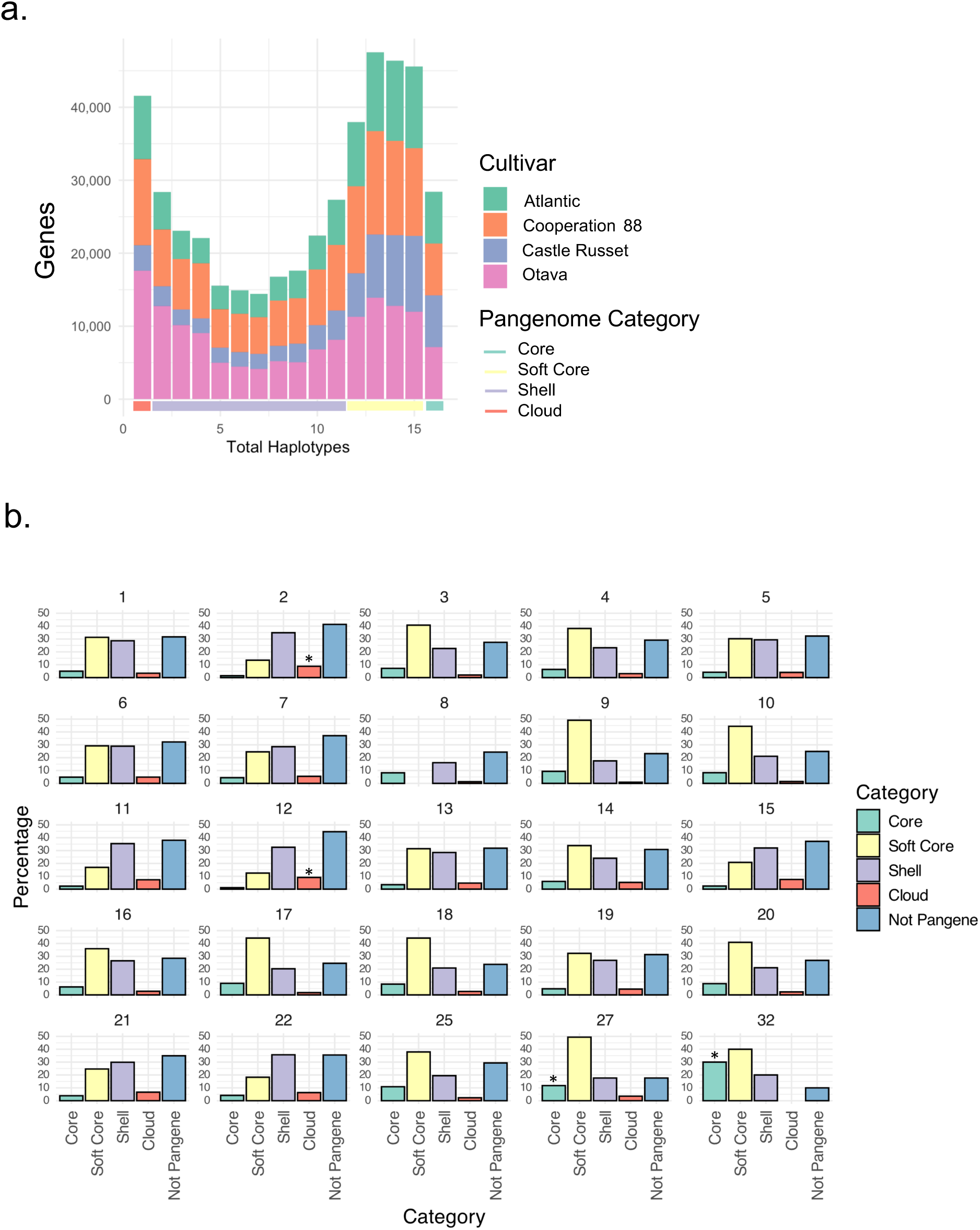
Pangenome Syntelogs and Their Module Membership. a. Distribution of pangenome syntelogs by the number of haplotypes in the syntelog group. The colors indicate the cultivar. The color bar on the x-axis indicates the category the haplotype belongs to: core genes include all 16 haplotypes, soft core genes 15-12 haplotypes, shell genes 2-11 haplotypes, and cloud genes are unique to individual cultivars. b. Percentage of genes in modules belonging to core, soft core, shell, cloud, and not part of the pangenome. Statistically significant membership to core, soft core, shell, and cloud are indicated with a star.

Atlantic is known for having an earlier bulking stage relative to the other cultivars, therefore we investigated the curated list of 218 genes involved in tuber initiation, growth and development for their membership in the core, soft core, shell, and cloud pangenome groups. Of the 218 genes, none were cloud genes suggesting that key genes involved in tuberization are conserved across cultivated potato. The core and soft core genes across all cultivars include 162 of the 218 genes and include *StSP6A* (tuberigen), the transcription factors: *StBRC1b/IT1, StPOTH1, StCO1/2*, and the histone modifier *StE(z)2*. The shell genes include transcription factors (*StBEL5, StBEL14,* and *StBEL22*), auxin transporters (*StPIN1* and *StPIN4*), and the sucrose transporter *StSWEET1*.

The gene coexpression modules were also examined for the enrichment of core, soft core, shell, and cloud genes to gain insights into conserved and diverged genes within cultivated potato (Figure 5b). We found that Modules 32 and 27 are enriched in core genes based on the hyper-geometric test enrichment value greater than 1 and a p-value less than 0.05 (Figure 5b). These modules have peak expression in stolon (Module 32) and tuber (Module 27) consistent with key roles of conserved genes in development. Module 32 has a functional annotation of stolon hormone regulation due to including the transcription factor *BRASSINAZOLE RESISTANT 1 (BZR1)* responsible for brassinosteroid synthesis, ABA signaling through *VP1/ABI3-LIKE 1*, and auxin related development through *indole-3-acetic acid inducible 26* (Overvoorde *et al*., 2005; Xu *et al*., 2025; Suzuki *et al*., 2007). Interestingly, in *A. thaliana BZR1* was found to be involved in the regulation of flowering by repressing *CO* (Xu *et al*., 2025). Orthologs of *BZR1* in potato could play a similar role in their regulation of *CO* which would ultimately affect tuberization through the *CO* role in regulating *SP5G*. This potential mechanism of *BZR1* could be conserved across cultivated potato as all three alleles were core.

### Enrichment of domestication genes

Domestication and improvement of potato reduced tuber glycoalkaloid content, allowed for adaptation to varying photoperiods, and reduced sexual fertility as the source-sink balance was shifted from sexual to asexual reproduction enhancing tuber size and yield (Lian *et al*., 2025; Hardigan *et al*., 2017; Pham *et al*., 2017). In particular, potato domestication involved genes that function in tuberization including *StCDF1* and *StSP6A* as well as a transcription factors involved in steroidal glycoalkaloid synthesis *GLYCOALKALOID METABOLISM 9* (*StGAME9*) (Hardigan *et al*., 2017; Lian *et al*., 2025). Previous studies focused on understanding domestication within potato identified 2,899 genes in the doubled monoploid reference accession *S. tuberosum* DM involved in domestication; these are syntenic with 12,065 alleles in Atlantic (Lian *et al*., 2025). Two pangenome categories, the core (p-value 3.07E-22) and soft core (p-value 4.59E-06), were significantly enriched in the genes under selection during domestication consistent with the cultivated status of these genomes that have undergone both domestication and selection for favorable traits.

Genes implicated in domestication (11,927 total) were enriched in Modules 3, 4, 6, 8, 9, 16, 17, 18, 20, 25, and 32 (p-value < 0.05). These represent modules with peak expression and hub gene-informed functional annotation in flower development and gene regulation (Modules 6 and 18), fruit development and gene regulation (Modules 9 and 20), leaf photosynthesis (Module 3), root gene regulation and response to stress (Module 4), tuber RNA binding and regulation of function (Modules 8 and 17), and stolon transcription factor activity and hormone regulation (Modules 16 and 32). As expected, these modules span development and regulation of gene expression. In both tomato and maize, domestication involved modification of transcriptional networks (Koenig *et al*., 2013; Swanson-Wagner *et al*., 2012) and similarly, a potato domestication study identified a set of transcription factors under selection either during domestication and/or subsequent improvement (Hardigan *et al*., 2017). We examined Submodule 16_3 that corresponds to tuber initiation and is enriched with genes involved in transcription regulation. In Submodule 16_3, domestication and/or improvement domestication genes are enriched with 97 genes (177 alleles) (p-value: 0.038) associated, including 13 transcription factors (23 genes) from the AP2/ERF-ERF, bHLH, C2C2, HB-HD-ZIP, LOB, MADS-MIK, MYB, OFP, and SRS families, suggesting modification of gene expression and coexpression network topology were selected by historical and modern potato breeders.

### Wild species introgressions are associated with stress responses

Wild species introgressions are known to contribute genes for disease, drought, and stress resilience (Hardigan *et al*., 2017; Lian *et al*., 2025) that are hypothesized to provide an adaptive function. A substantial proportion of stress-responsive genes in the expression atlas were derived from wild species introgressions, with 28.3–35.9% of DEGs under stress conditions. The stress-responsive conditions which were considered enriched in wild species introgression include drought root, drought leaf (downregulated, p-value: 1.04E-6), and MeJa treated leaf (upregulated, p-value: 0.001). For example, drought root upregulated and downregulated DEGs were enriched for wild species introgressions; of the 4,910 upregulated genes, 1,620 were derived from wild species introgressions (p-value: 0.004). Additionally, of the 8,293 downregulated genes, 2,754 are from wild species introgressions (p-value: 3.61E-5). The DEGs from wild introgressions are distributed across all 25 modules. However, no modules were enriched for wild species introgression. Stress responses can be associated with particular modules through their functional annotation and peak expression. For example, Module 21 was functionally annotated as “root transcriptional regulation in response to drought” and had peak expression during drought treatment in roots (Figure 3a). Module 21 contains 218 hub genes, including 24 transcription factors, 10 of which originate from wild species introgressions. These transcription factors belong to key families associated with stress responses, including AP2/ERF-ERF, HB-HD-ZIP, HSF, MYB, and NAC. NAC transcription factors were previously identified as central to drought response in potato (Wen *et al*., 2025). Notably, NAC transcription factors are also well-documented contributors to drought resilience in multiple crop species, such as GhNAC3 in cotton, JUNGBRUNNEN1 in tomato, TaNAC071-A in wheat, and STRESS-RESPONSIVE NAC 1 in rice (Hu et al., 2006; Thirumalaikumar et al., 2018; Mao et al., 2022; Xia et al., 2023). In wheat, the NAC has increased expression due to an insertion that contains MYB cis-regulatory elements (Mao *et al*., 2022). In Module 21 there are three wild species introgressed MYB transcription factors that could potentially exhibit a similar function given their placement within the same coexpression module. These findings highlight the critical role of wild species introgression in enhancing drought tolerance, supported by both the expression patterns observed and functional parallels in other plant systems.

## CONCLUSION

A comprehensive, allele-resolved 34 tissue and treatment atlas in the tetraploid potato cultivar Atlantic has provided valuable insights into tuber development and stress responses. Creation of a robust gene coexpression network revealed 26 modules that were categorized based on their peak expression and hub gene GO term annotations. Structural variation across four unique cultivars established the core, soft core, shell, and cloud genes from a pan-genomic analysis and revealed modules enriched in core and cloud genes providing insights into conserved and Atlantic specific genes. Domestication and wild species introgression data provided deeper insights into gene coexpression networks identifying genes necessary for tuberization and sources of genetic diversity. These findings offer a valuable resource for future research and breeding efforts aimed at understanding tuberization and improving potato resilience and yield.

## MATERIALS AND METHODS

### Plant growth & tissue harvest

*Solanum tuberosum* cv Atlantic were propagated from tissue culture plantlets and transferred to a growth chamber for growth and subsequent sampling. Tissues were collected from a stolon/tuber developmental series, diurnal time course (leaves), flowers, fruits, stems, roots, and a set of abiotic/biotic stress conditions. For the leaf time-course, sampling of mature plants occurred every four hours after starting at 6am (dawn) for 24 hours. Plants were maintained in a controlled growth chamber with day/night temperatures of 25.5 °C/15.5 °C and a 15-hour photoperiod. For Methyl Jasmonate (MeJA) treatment, 250µM MeJA was applied with a spray bottle until leaves were drenched, samples were collected 24 hours after application. For benzothiadiazole (BTH) treatment, a 100µg/ml solution of BTH was applied via a spray bottle until leaves were drenched followed by sampling 24 hours after treatment. For salt treatment, 100mL of a 150mM solution of NaCl was applied to each pot and samples were collected 24 hours later. For cold treatment, conditions were changed to 10°C constant and samples collected after 24 hours of treatment. For heat treatment, plants were watered to avoid drought stress prior to increasing the temperature to 37°C (day) and 28°C (night); samples were collected after 24 hours. For drought treatment, water was withheld until visible wilting was observed after which plants were sampled (seven days).

### RNA isolation, library construction, and sequencing

RNA was isolated using a modified hot borate protocol (Wan & Wilkins, 1994). TURBO DNase (Invitrogen, Waltham, MA) was used to remove any residual DNA from all samples. RNA-Seq libraries were generated using the Illumina Stranded mRNA Prep Ligation kit with IDT for Illumina RNA Unique Dual Indexes and sequenced on a NovaSeq 6000 in paired end mode to 50 nt in length (Illumina, San Diego, CA). Sequencing was performed by Texas A&M AgriLife Research: Genomics and Bioinformatics Service.

### Expression abundance estimation and tissue specific expression

Illumina RNA-Seq reads were evaluated for quality using FastQC (https://www.bioinformatics.babraham.ac.uk/projects/fastqc/) and MultiQC (Ewels *et al*., 2016). Reads were cleaned using Cutadapt (v4.2) (Martin, 2011) with a 3’ adapter provided for both read one and read two, a minimum read length (-m) of 40 nt, low quality reads trimmed from the 3’ end of each read (-q 30), flanking N’s removed (--trim-n), and the number of times to trim an adapter set to two (--times 2). Reads were checked for contamination using Kraken2 (v2.1.2) (Wood *et al*., 2019) with the database k2_pluspfp_20220908. Based on the median total reads after cleaning, 43 million reads were subsampled using seqtk (v1.3; https://github.com/lh3/seqtk). To determine expression abundances, cleaned subsampled reads were pseudoaligned to the Atlantic v3 (Hoopes *et al*., 2022) high confidence representative gene set using kallisto quant (v0.4.8) (Bray *et al*., 2016) with the --rf-stranded option and the k-mer size set to 21. Genes were removed from downstream analysis if they had a TPM of zero across all libraries. Libraries were flagged based on GC content, percent of reads discarded during cleaning, over-represented sequences, contamination, pseudoalignment rate, and Pearson correlation coefficient between biological replicates. Libraries with multiple flags were evaluated individually and discarded based on results.

Tissue specific expression analysis was conducted using the tspex with default parameters (Camargo *et al*., 2020) (v.0.6.2). Allele groups were determined between *S. tuberosum* DM (v.6.1) (Pham *et al*., 2020) and Atlantic using GENESPACE (Lovell *et al*., 2022) (v1.2.3) with the representative high confidence model proteins for each haplotype and the expected ploidy set to one. Preferential allele expression was determined by performing a t-test within the allele groups. A preferentially expressed allele had a log2 fold change value > 2 and false discovery rate correction < 0.05.

### Differentially expressed genes

Differentially expressed genes were identified between the abiotic/biotic stress treatments and their controls using the edgeR package in R (Chen *et al*., 2025) (v.3.42.4). Salt treatment control was the proper control for the drought and salt treatments. Genes were considered differentially expressed if they have a log fold change greater than 2 and p-value less than 0.05.

### Coexpression network construction

Gene coexpression networks were generated from high confidence representative models. The networks were constructed using Simple Tidy GeneCoEx (Li *et al*., 2023) where all gene models were used in the coexpression networks. Networks were visualized using Cytoscape (Shannon *et al*., 2003) (v.3.10.1). Hub genes were determined by calculating the number of edges connected to nodes and categorizing the top 5% of the genes in the module as hub genes.

### Gene ontology term enrichment analysis

Atlantic_v3 GO terms were obtained from SpudDB (https://spuddb.uga.edu/) and enrichment analysis was conducted using the topGO package in R with the algorithm set to “weight01” and the statistic specified as “fisher” (Alexa and Rahnenfuhrer, 2024; R Core Team, 2023) (v.2.52.0). Terms with a p-value less than 0.05 were considered significant. GO term annotations were mapped using the GO.db package in R (Carlson, 2023; R Core Team, 2023)(v.3.17.0). *StSP6A* submodule GO terms were placed into categories based on their GO term ancestors.

### Comparative genome analyses

Syntelogs were identified within Atlantic v3 (Hoopes *et al*., 2022), Castle Russet (Hoopes *et al*., 2022), Cooperation 88 (Bao *et al*., 2022), and Otava (Sun *et al*., 2022)using GENESPACE (Lovell *et al*., 2022) (v1.2.3) with the representative high confidence model proteins for each haplotype and setting the expected ploidy to one. The syntelog sets were extracted from the genes denoted as PASS in the pangenome database file. The genes were then binned into pangenome categories based on the number of haplotypes present in the syntelog sets where 1 haplotype is a cloud gene, 2-11 haplotypes is a shell gene, 12-15 haplotypes is a soft core gene, and 16 haplotypes is a core gene. Enrichment analysis for pangenome categories in coexpression modules were performed using the phyper function in R (R Core Team, 2023).

### Selection during domestication and wild species introgression enrichment analysis

Selection during domestication and wild species introgression datasets were obtained from Lian et al., 2025. The genes from these datasets were from *S. tuberosum* DM (v.6.1) and were mapped to pangenome syntelogs to identify corresponding genes in Atlantic. Enrichment analysis for the domestication pressure and wild species genes in modules was conducted using the phyper function in R (R Core Team, 2023).

## DATA AVAILABILITY

Raw sequences are available in the National Center for Biotechnology Information Sequence Read Archive under BioProject PRJNA753086.

## ACKNOWLEDGEMENTS

We are thankful for funding from the National Science Foundation (IOS-2140176) to C.R.B., Michigan State University to J.B., National Institutes of Health T32 Training Grant to D.M. (5T32GM142623), University of Georgia (C.R.B.), Georgia Research Alliance (C.R.B.), and Georgia Seed Development (C.R.B.). We especially thank Kennedy Baldwin, Caleb Fisher, and Rachel Shereda for RNA isolation and tissue harvest.

